# E2F1 Drives Endothelial Arterial Programming in Pulmonary Arterial Hypertension

**DOI:** 10.64898/2026.07.02.736230

**Authors:** Dan Yi, Ankit Tripathi, Qi Zheng, Bin Liu, Susie Weizhi Cao, Jeffrey R. Koenitzer, Mengcheng Shen, Michael B. Fallon, Zhiyu Dai

## Abstract

**Background:** Pulmonary arterial hypertension (PAH) is driven by maladaptive endothelial remodeling, but the transcriptional regulators that couple proliferative stress to arterialized endothelial states remain incompletely defined. E2F transcription factor 1 (E2F1) is classically viewed as a cell-cycle regulator; whether E2F1 functions as a disease-driving node that promotes endothelial arterial programming in PAH remains unknown.

**Methods:** We integrated human PAH lung transcriptomic analyses, deconvolution-based endothelial-state scoring, and complementary mouse and rat PH models with bulk RNA-seq, single-cell RNA-seq, pseudotime analysis, and CellChat inference. E2F1 function was tested using adenoviral E2F1 overexpression, pharmacological pan-E2F inhibition with HLM006474, and genetic *E2f1* loss on a tamoxifen-inducible endothelial *Egln1*-deletion background.

**Results:** In PAH lungs, E2F1 was increased and arterial endothelial cell (AEC) fraction and expanded arterial program scores were elevated. Similarly, *Egln1^Tie2Cre^* lungs showed increased E2F1, induction of arterial remodeling genes, and activation of an E2F target program. Genetic loss of *E2f1* reduced right ventricle systolic pressure, right ventricle hypertrophy, vascular remodeling, and distal muscularization in *Egln1*-driven PH mice model. Bulk RNA-seq showed suppression of E2F, mitotic, epithelial mesenchymal transition, and extracellular matrix-remodeling programs. Single-cell RNA-seq showed reduced AEC accumulation, normalized CAP1/CAP2 distribution, and reduced progression along the CAP1-AEC trajectory. CellChat analysis identified loss of an arterial communication hub, including reduced ECM, VEGF, and Notch signaling when E2F1 is loss. Conversely, E2F1 overexpression in human lung microvascular ECs increased proliferation, activated E2F/cell-cycle and Notch/arterial programs. Pharmacological inhibition of E2F via HLM006474 suppressed endothelial proliferation and attenuated *Egln1*-driven and MCT-induced PH, including reversal of established MCT-PH.

**Conclusions:** E2F1 acts as a disease-relevant transcriptional factor linking endothelial cell-cycle activation to arterial programming, matrix and angiogenic communication programs, and pulmonary vascular remodeling. Genetic or pharmacological E2F inhibition mitigates experimental PH, supporting E2F1 as a therapeutic target in PAH.

**Clinical Perspective What is new?:** 1. This study identifies E2F1 as a previously unrecognized driver of PAH rather than only a downstream marker of cell-cycle activation.

2. Genetic loss of E2f1 rescues hemodynamic and structural features of Egln1-driven PAH, and pharmacological E2F inhibition attenuates both Egln1-driven and monocrotaline-induced PH.

3. Mechanistically, E2F1 links endothelial proliferation to Notch-associated arterial programming, AEC accumulation, and CAP1-to-iAEC-to-AEC trajectory progression.

**What are the clinical implications?:** 1. E2F1 defines a tractable transcriptional node that integrates proliferative stress with arterial endothelial reprogramming, a core pathological feature of PAH vascular remodeling.

2. Pan-E2F small-molecule inhibitors, several of which are in development for oncology, may be repurposable for PAH if E2F1-dependent endothelial arterial-programming signatures identify responsive disease states.

3. Plasma- or tissue-based readouts of E2F1 activity may identify PAH patients most likely to benefit from E2F-directed therapy.

## Introduction

Pulmonary arterial hypertension (PAH) is a progressive disease characterized by occlusive remodeling of small pulmonary arteries, sustained elevation of pulmonary vascular resistance, right ventricular failure, and premature death.^1–3^ Although therapies targeting nitric oxide, endothelin, and prostacyclin signaling improve symptoms and outcomes, PAH remains incurable, particularly in idiopathic and heritable disease.^1,4^ The core histopathological lesion reflects coordinated reprogramming of pulmonary endothelial cells (ECs) toward apoptosis-resistant, hyperproliferative, inflammatory, and remodeling-permissive states. Defining the transcriptional circuits that stabilize this maladaptive endothelium is therefore essential for developing disease-modifying therapy.

Dysregulated transcription factor activity is a central feature of pulmonary endothelial pathobiology in PAH. Loss-of-function variants in BMPR2 and SOX17, gain of HIF-2α activity, and altered FOXM1, KLF2, KLF4, NRF2, STAT3, and YAP/TAZ signaling converge on the proliferative and apoptosis-resistant EC phenotype that defines pulmonary vascular lesions.^5–10^ Among proliferation-linked transcription factors, the E2F family is uniquely positioned downstream of the retinoblastoma-cyclin/CDK axis to license S-phase entry, DNA replication, mitotic progression, and nucleotide metabolism.^11–13^ E2F1 can also engage tumor-suppressive programs through CDKN2A/ARF, p53 stabilization, and CDKN1A/p21 induction, coupling proliferative stress to apoptosis or senescence depending on context.^14–16^

In the pulmonary circulation, E2F1 has been studied mainly in smooth muscle and only recently in endothelial dysfunction. Na+/H+ exchanger-dependent downregulation of p27 and activation of E2F1 promotes pulmonary arterial smooth muscle cell migration and proliferation,^17^ and our prior work identified E2F1 as a downstream effector by which endothelial SOX17 loss drives EC dysfunction and PH.^18^ Recent lineage-tracing and single-cell studies further show that capillary ECs can transition toward an arterial state in PH through a HIF-2α/Notch4 axis.^19^ However, whether E2F1 is itself required for PAH progression in vivo, and whether E2F1 controls this pathological endothelial arterial-programming process rather than only cell-cycle entry, remain unknown.

We therefore hypothesized that E2F1 upregulation in pulmonary ECs is a driver, rather than a bystander, of pulmonary vascular remodeling and PAH. Here we show that E2F1 is induced in human PAH samples and in multiple preclinical PH models; that loss of E2F1 attenuates *Egln1*-driven PH in vivo; that E2F1 gain of function in HLMVECs promotes proliferation, suppresses BMPR2 signaling, activates Notch programs, and induces an arterial-programming signature; and that pharmacological E2F inhibition with HLM006474 attenuates experimental PH. These findings establish E2F1 as a novel PAH driver and reveal an unexpected mechanism by which E2F1 couples cell-cycle activation to maladaptive endothelial arterial programming.

## Methods

The data, analytic methods, and study materials that support the findings of this study will be made available from the corresponding author upon reasonable request, to the extent allowed by institutional policy. Bulk and single-cell RNA-sequencing data have been deposited in the Gene Expression Omnibus under accession numbers GSE335785 and GSE335786, respectively. The scripts to generate the data have been deposited in the GitHub (DaiZYlab).

### Human samples

De-identified archived human lung tissue from patients with idiopathic PAH (IPAH) and unused donor/failed-donor controls were obtained from the Pulmonary Hypertension Breakthrough Initiative (PHBI) under approved institutional protocols, as described previously.^18,20^ De-identified archived lung sections from IPAH and control donors were used for RNAScope analysis. All human tissue were approved by the Institutional Review Boards of Washington University in St. Louis and/or the University of Arizona, and all samples were handled without direct subject identifiers. Human lung bulk RNA-seq analyses used the GSE254617 cohort for failed-donor (FD; n = 52) and IPAH (n = 40) comparisons. Differential expression was modeled with DESeq2^21^ using a design that included sequencing batch and disease group. Deconvolution-estimated AEC fractions were obtained from LungMAP-reference cell-type deconvolution at the subject level. The expanded arterial program score was calculated from 19 genes (BMX, CXCL12, DLL4, EFNB2, GJA5, HEY1, HEY2, JAG1, NOTCH4, NRP1, SOX17, APOLD1, DEPP1, EDN1, GNG11, IGFBP7, LTBP4, TMEM88, and UACA) as the mean of gene-wise z-scored log2-normalized expression values and then centered to the mean FD score for display. Human score and deconvolution comparisons were tested with two-sided Welch’s t tests, with Mann-Whitney testing used as a sensitivity analysis.

### Animals

All animal procedures were approved by the Institutional Animal Care and Use Committees of Washington University in St. Louis and/or the University of Arizona and were performed in accordance with NIH guidelines and the Guide for the Care and Use of Laboratory Animals. *Egln1^Tie2Cre^*, *Egln1^Cdh5CreERT^*^2^ mice have been described previously and maintained in the Dai lab.^22,23^ *E2f1^-/-^* mice were obtained from JAX (#002785).^24^ Both male and female mice were included; sex was recorded as a biological variable and balanced across experimental groups when possible.

Tamoxifen-inducible endothelial *Egln1* deletion was generated by crossing *Egln1^flox/flox^* mice with Cdh5-CreERT2 mice (iCKO). *E2f1*^−/−^;*Egln1^Cdh5CreERT^*^2^ mice (iDKO) were generated by intercrossing *E2f1^−/−^* mice with the iCKO line. Cre recombination in Cdh5-CreERT2 cohorts was induced at 6–8 weeks of age by intraperitoneal tamoxifen (100 mg/kg/day for 5 consecutive days; Sigma T5648, dissolved in corn oil), and mice were analyzed 6–8 weeks after the final tamoxifen dose unless otherwise specified.

For hypoxic exposure studies, mice were housed in normobaric hypoxia (10% O_2_) for 3 weeks in a ventilated hypoxic chamber, while control mice were maintained in room air (21% O2). Littermate controls were used when available.

### Monocrotaline (MCT) rat model

Adult male Sprague-Dawley rats (6 weeks old, 150–160 g) received a single subcutaneous injection of monocrotaline (MCT, 33 mg/kg; Cayman) to induce PH development.^25^ For E2F inhibition, rats received HLM006474 (25 mg/kg, intraperitoneal; Sigma 324461) or vehicle (Saline) five times a week. In prevention studies, dosing began 2 days after MCT injection and continued until terminal hemodynamic assessment. For reversal studies, treatment was initiated 2 weeks after MCT injection and continued for 2 weeks.

### Pharmacological E2F1 inhibition in *Egln1^Cdh5CreERT^*^2^ mice

Tamoxifen-induced *Egln1^Cdh5CreERT^*^2^ (iCKO) mice were randomized to vehicle or HLM006474 treatment beginning 2 weeks after the final tamoxifen dose. HLM006474, a small-molecule E2F inhibitor,^26^ was administered intraperitoneally at 25 mg/kg three times per week for 4 weeks. Mice underwent terminal hemodynamic assessment and tissue collection at the end of treatment.

### Hemodynamics and right ventricular remodeling

Right ventricular systolic pressure (RVSP) was measured by catheterization of the right ventricle using a Millar Mikro-tip pressure catheter (SPR-1000 for mice and SPR-869 or equivalent for rats), as described in prior studies.^22,27^ Animals were anesthetized with 1.5–2% isoflurane for the present experiments unless otherwise indicated, and pressure waveforms were recorded after stable catheter positioning. Following RVSP measurement, hearts were excised and the right ventricular free wall (RV) was dissected from the left ventricle plus interventricular septum (LV+S). Right ventricular hypertrophy was calculated as RV/(LV+S) (Fulton index).

### Histology, immunofluorescence and RNAScope

Lungs were perfused free of blood, inflation-fixed with 4% paraformaldehyde at constant pressure, paraffin-embedded or cryo-embedded as appropriate, and sectioned for histology and immunostaining. Russell-Movat pentachrome and α-smooth muscle actin (α-SMA; Abcam #ab5694) staining were used to assess pulmonary vascular remodeling, medial wall thickening and distal muscularization. α-SMA^+^ muscularization was quantified in distal vessels by size class (<50 μm and >50 μm), using multiple random fields per animal, consistent with prior lab morphometry. Wall thickness on Russell-Movat-stained sections was quantified by normalizing medial wall thickness to vessel diameter/radius as in prior publications.

For RNAScope, sections were incubated with E2F1 probes (ACD, #592801) counterstained with DAPI according to the vendor’s protocol, and imaged by confocal microscopy.

### Adenoviral E2F1 overexpression in HLMVECs

Primary human lung microvascular endothelial cells (HLMVECs; Lonza CC-2527) were cultured in EBM-2 medium supplemented with EGM-2 MV components and used at passages 4–7. Cells were transduced with adenovirus expressing human E2F1 (AdvE2F1, SignaGen Laboratories) or control adenovirus expressing GFP (Ad-GFP/AdvCTL) at MOI 5 in basal medium for 24 h, followed by recovery in complete medium before assays. E2F1 induction was confirmed by qPCR and immunoblotting before functional or transcriptomic analyses.

### siRNA knockdown of E2F1

For E2F1 knockdown, HLMVECs were transfected with human E2F1 siRNA (Santa Cruz Biotechnology, sc-29297) or control siRNA (Santa Cruz Biotechnology, sc-37007) using the HMVEC-L Nucleofector Kit (Lonza, VPB-1003) and Nucleofector 2b device (Lonza, AAB-1001; program A-034), consistent with the published SOX17/E2F1 protocol. Based on the published method, siRNA was used at 50 nM unless otherwise indicated. Cells were harvested 72 h after transfection for RNA anda protein. Knockdown efficiency was confirmed by qPCR and immunoblotting.

### Proliferation assays

Endothelial proliferation was measured by BrdU incorporation followed by BrdU/DAPI immunofluorescence. For AdvE2F1 experiments, HLMVECs were infected with Adv-E2F1 or Adv-CTL (MOI = 5) for 48 hours and pulsed with BrdU (10 μM, 4 h) before fixation and quantification of BrdU-positive nuclei. For pharmacological inhibition studies, HLMVECs were treated with HLM006474 (40 μM) or vehicle under VEGF-A stimulation (0.5 nM) or hypoxia (10% O2) overnight, and BrdU incorporation was quantified from multiple fields per condition.

### Bulk and single-cell RNA sequencing

Bulk RNA-seq was performed on HLMVECs transduced with AdvE2F1 or control adenovirus (Ad-GFP; AdvCTL) (n = 3/group, MOI = 5, 48 hours), HLMVECs transfected with si-E2F1 or si-control (n = 3/group), and whole lungs from tamoxifen-induced iDKO and iCKO mice (n = 4/group). Libraries were prepared using the Zymo-Seq SwitchFree 3′ mRNA Library Kit (Zymo Research #R3009) and sequenced on an Illumina platform to a target depth of approximately 30 million paired-end reads per sample. Reads were aligned to GRCh38 (human) or GRCm39 (mouse), gene-level counts were analyzed with DESeq2, and P values were adjusted by the Benjamini-Hochberg method. Genes with |log2FC| > 1 and adjusted P < 0.05 were considered significantly differentially expressed unless otherwise specified. Heatmaps used variance-stabilized or normalized expression values with row-wise scaling as indicated.

Single-cell RNA-seq was performed on whole-lung cells from iCKO and iDKO mice to profile global lung cell composition. Single-cell libraries were generated using the 10X Genomics fixed RNA Profiling Reagent Kits for Multiplexed Samples (PN-1000414) and Chromium platform, sequenced on an Illumina instrument to a target depth of approximately 20,000 reads per cell, and processed with Cell Ranger. Downstream analyses were performed in Seurat after integration across genotypes. The final annotated dataset contained 16,978 cells, including 9,364 iCKO cells and 7,614 iDKO cells, across 31 manually curated cell populations. Cluster annotation used automated label transfer and manual curation supported by published lung/endothelial atlases and LungMAP CellCards marker genes^28^. Cell-type-specific differential expression used Seurat FindMarkers with the Wilcoxon rank-sum test (ident.1 = iDKO, ident.2 = iCKO, logfc.threshold = 0.25, min.pct = 0.1), followed by adjusted P-value correction. Cell-type proportion changes were assessed with scProportionTest permutation testing. Endothelial cells were computationally subsetted for EC subtype, trajectory, pseudotime, and differential-expression analyses.

Gene set enrichment analysis (GSEA) was performed against MSigDB HALLMARK and KEGG gene sets using clusterProfiler.^29^ Genes were ranked by log2 fold change for the relevant comparison, and pathway significance was determined by permutation-based testing with false-discovery-rate correction. For over-representation analysis, differentially expressed genes were used as input, and adjusted P < 0.05 was considered significant unless otherwise indicated.

Cell-cell communication was inferred with CellChat v2.1.2^30^ using the mouse ligand-receptor database. Condition-specific CellChat objects were built from log-normalized RNA assay data across all 31 annotated cell types, with communication probabilities computed using triMean and population-size correction. Interactions were filtered at a minimum of 10 cells per group, and iCKO and iDKO CellChat objects were merged for differential interaction, information-flow, pathway, and ligand-receptor analyses.

For endothelial trajectory analysis, Slingshot was run on the EC UMAP using CAP1 as the root state and EC subtype labels CAP1, CAP2, iAEC, AEC, and VEC. The disease-relevant trajectory was defined as the CAP1-iAEC-AEC lineage. Dynamic genes along this lineage were identified with tradeSeq fitGAM using four knots after filtering genes detected in at least 20 cells with total raw counts ≥200; associationTest P values were adjusted by the Benjamini-Hochberg method, and FDR < 0.05 was considered significant.

### qPCR and immunoblotting

Total RNA was isolated using the Zymo Quick-RNA Miniprep kit, reverse-transcribed with Applied Biosystems™ High-Capacity cDNA Reverse Transcription Kit and analyzed by SYBR Green qPCR. Relative gene expression was calculated using the comparative Ct method and normalized to housekeeping controls (primers for human *E2F1*: CCGGGGAATGAAGGTGAACA, GAGCAAAAGGGCCGAAAGTG; primers for human 18S rRNA: TTCCGACCATAAACGATGCCGA, GACTTTGGTTTCCCGGAAGCTG).

Whole-cell or tissue lysates were prepared in RIPA buffer containing protease/phosphatase inhibitors and analyzed by SDS-PAGE/immunoblotting. Primary antibodies included E2F1 (CST 3742), BMPR2 (BD 612292), SOX17 (CST 81778) and β-actin (Sigma A5441), with HRP-conjugated secondary antibodies and ECL detection.

## Statistical analysis

Data are presented as mean ± SD unless otherwise indicated. Two-group comparisons were analyzed by unpaired two-tailed Student’s t test, with Welch’s correction when variances were unequal. Multiple-group comparisons were analyzed by one way ANOVA with Tukey post hoc correction. Survival curves were compared by log-rank testing. For sequencing analyses, differential expression P values were adjusted with the Benjamini-Hochberg false-discovery-rate method, and pathway analyses used the FDR thresholds specified in each analysis. A two-sided P < 0.05 was considered statistically significant. Analyses were performed in GraphPad Prism and R.

## Results

### E2F1 dysregulation accompanies arterial endothelial remodeling and E2F target activation in PAH

Our previous studies demonstrated that E2F1 activation mediates SOX17 deficiency-induced endothelial dysfunction and PH development^18^. To assess *E2F1* dysregulation in human PAH, we reanalyzed lung bulk RNA-seq from patients with idiopathic PAH (IPAH) and failed-donor (FD) controls. *E2F1* mRNA was modestly higher in IPAH lungs, although the difference was not statistically significant (P = 0.1206; **Figure 1A**). Representative RNAScope imaging showed stronger *E2F1* staining in IPAH than control lungs (**Figure 1B and 1C**).

**Figure 1.**
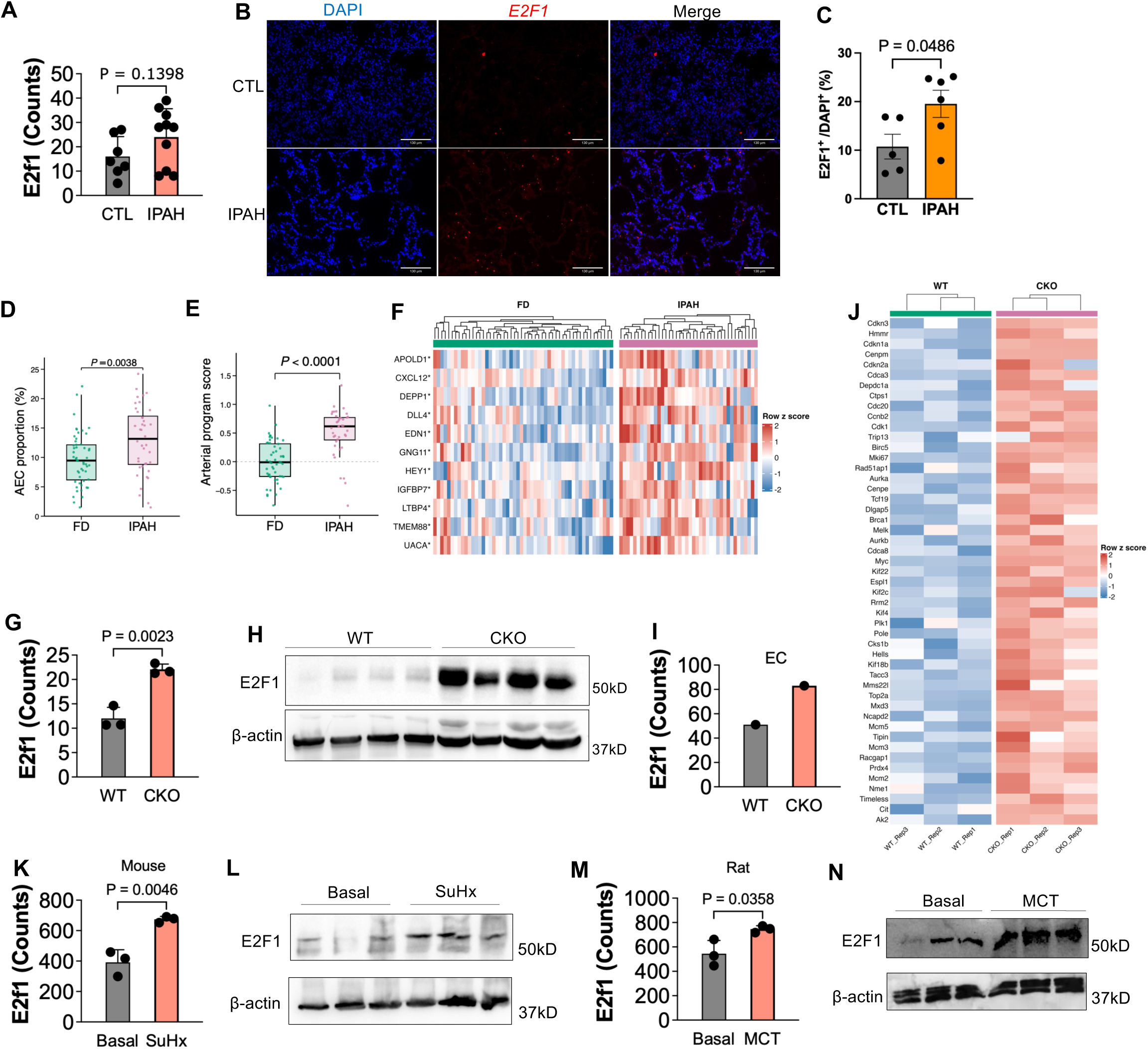
E2F1 is upregulated in pulmonary tissues from IPAH patients and in preclinical PH models. (A) Bar graph showing E2F1 mRNA expression from bulk RNA-seq of lung tissues from healthy controls (Control, n = 7) and idiopathic PAH (IPAH) patients (n = 10). Data are presented as normalized counts. (B and C) Representative RNAScope images and quantification of lung sections from a healthy donor (CTL) and an IPAH patient. Sections were stained with DAPI (blue, nuclei) and E2F1 probes (red). Scale bar = 130 µm. (D) Deconvolution-estimated arterial endothelial cell (AEC) proportion in failed-donor controls (FD; n = 52) and patients with IPAH (n = 40). (E) Expanded arterial program score calculated as the mean standardized expression of 19 arterial-associated genes and centered to the mean FD score. In D and E, points represent individual samples; boxes indicate the median and interquartile range, and whiskers extend to 1.5 times the interquartile range. P values were determined using two-sided Welch’s t tests. (F) Heatmap showing the 11 arterial genes significantly upregulated in IPAH lungs (DESeq2 FDR & 0.05 and log2FC > 0). (G) E2F1 mRNA expression in whole-lung lysates from wild-type (WT) and Egln1Tie2Cre (CKO) mice. (H) Representative immunoblots of E2F1 and β-actin. (I) E2F1 mRNA expression in isolated lung ECs from CKO mice. (J) Heatmap showing row-scaled expression of E2F target genes in WT (n = 3) and CKO (n = 3) mouse lungs. Color scale indicates row z score of log2-transformed normalized counts (blue: lower expression; red: higher expression). (K and L) E2F1 mRNA and protein levels in SuHx mouse lungs compared with basal controls. (M and N) E2F1 mRNA and protein expression in whole-lung lysates from vehicle-treated (Basal) or monocrotaline (MCT)-treated rats at 3 weeks post-injection. For animal and cell experiments, individual points indicate biological replicates; two-group comparisons used unpaired two-tailed t tests unless otherwise indicated.

Our previous data demonstrated the arterial reprogramming in preclinical and human PAH^19^. Because changes in endothelial composition can influence whole-lung transcriptomic profiles, we used deconvolution to estimate the arterial endothelial cell (AEC) fraction from a previously published bulk RNA-seq dataset from PAH patients^31^. The estimated AEC fraction was higher in IPAH than FD lungs (12.93% versus 9.66%; P = 0.0038; **Figure 1D**). An expanded 19-gene arterial program score was also increased in IPAH (IPAH-minus-FD mean difference; P < 0.0001; **Figure 1E**). Consistent with this score, a heatmap restricted to genes that were both upregulated and significant at FDR < 0.05 showed coordinated increases in arterial EC markers including *APOLD1*, *CXCL12*, *DEPP1*, *DLL4*, *EDN1*, *GNG11*, *HEY1*, *IGFBP7*, *LTBP4*, *TMEM88*, and *UACA* (**Figure 1F**).

We next examined E2F1 activation in *Egln1^Tie2Cre^* (CKO) mice, a severe PH mouse model^22^. Whole-lung E2F1 mRNA and protein were markedly increased in CKO compared with wild-type (WT) lungs (P = 0.0023; **Figure 1G, H**), and isolated lung ECs from CKO mice showed parallel E2F1 induction (**Figure 1I**). Because PAH-associated endothelial remodeling includes expansion of arterial-like endothelial states^19,32–34^, we examined a focused arterial remodeling gene panel in WT and CKO lungs. This heatmap showed coordinated induction of arterial genes in CKO lungs, including Cxcl12, Sox17, Apold1, Edn1, Gng11, Igfbp7, Tmem88, Uaca, Hey1, Eng, Col4a1, Col4a2, Sparc, and Hspg2 (**Supplemental Figure 1**). We also observed an induction of E2F target program in CKO lungs (**Figure 1J**). Together, these data indicate that E2F1 induction in *Egln1*-driven PH coincides with activation of an arterial/endothelial remodeling program and an E2F target program.

E2F1 induction was also observed in two additional experimental PH models.^35^ In mice exposed to SU5416 plus hypoxia (SuHx), lung E2F1 protein was higher than in basal controls (P = 0.0046; **Figure 1K, L**). Likewise, monocrotaline (MCT)-treated rats showed increased lung E2F1 mRNA and protein (P = 0.0358; **Figure 1M, N**). Together, these findings link E2F1 induction across experimental PH models and human PAH with an arterialized endothelial phenotype and E2F-associated proliferative program.

### Genetic deletion of *E2f1* inhibits *Egln1*-driven PH

To test whether E2F1 is required for PH development *in vivo*, we generated tamoxifen-inducible, endothelial-restricted *Egln1* deletion in an *E2f1*-deficient background (iDKO). The breeding strategy is shown in **Figure 2A**. Loss of E2F1 protein in iDKO and global *E2f1* knockout (KO) lungs was confirmed by immunoblot (**Figure 2B**). iCKO mice developed robust PH 6–8 weeks after tamoxifen, with significantly elevated RVSP and increased RV/(LV+S) ratio. Deletion of *E2f1* (iDKO) significantly reduced RVSP compared to iCKO littermates (**Figure 2C**). Consistently, iDKO mice also showed a significant reduction in RV/LV+S ratio (**Figure 2D**). Russell-Movat pentachrome staining showed extensive medial wall thickening in iCKO mice, which was attenuated in iDKO lungs (**Figure 2E, F**). α-SMA immunofluorescence revealed a marked increase in fully muscularized distal vessels in iCKO mice, and this muscularization was significantly reduced in iDKO mice (**Figure 2G, H**). These data establish that E2F1 is required for the development of *Egln1*-driven PH in mice.

**Figure 2.**
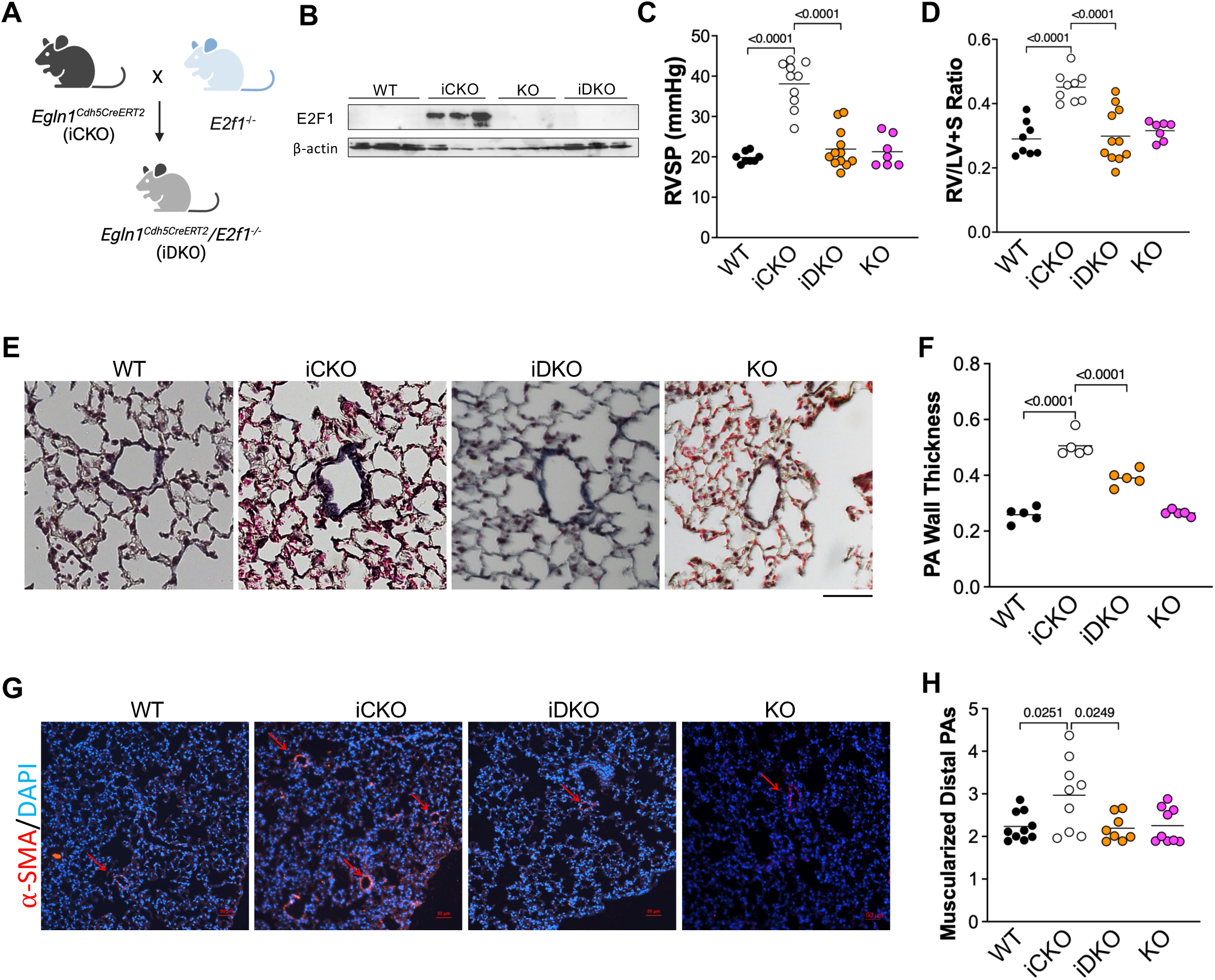
Genetic deletion of *E2f1* rescues *Egln1*-driven PH. (A) Breeding strategy for generating iDKO mice. iCKO (Egln1Cdh5CreERT2) mice were crossed with global E2f1 knockout mice (E2f1−/−, KO). (B) Representative immunoblots of E2F1 protein in whole-lung lysates from WT, iCKO, KO, and iDKO mice 6 weeks after tamoxifen induction. (C) Right ventricular systolic pressure (RVSP) in the indicated genotypes 6 weeks after tamoxifen induction. (D) Right ventricular hypertrophy quantified as Fulton index, RV/(LV+S). (E and F) Pulmonary vascular remodeling in WT, iCKO, KO, and iDKO mice. (E) Representative Russell-Movat pentachrome-stained distal pulmonary arteries (20-100 µm diameter). Scale bar = 50 µm. (F) Quantification of medial wall thickness in distal pulmonary arteries. (G and H) Pulmonary arterial muscularization. (G) Representative α-SMA/DAPI immunofluorescence images of distal pulmonary arteries. Scale bar = 50 µm. (H) Quantification of muscularized arteries in vessels <50 µm and >50 µm diameter. Individual points indicate animals, and n per group is shown in each graph. Multiple-group comparisons were analyzed by one-way ANOVA with Tukey’s post hoc test.

### Bulk RNA-seq shows E2f1 deletion suppresses cell-cycle and mitotic programs in *Egln1*-driven PH lungs

To define transcriptomic changes associated with the hemodynamic rescue observed in iDKO mice, we performed whole-lung bulk RNA-seq comparing tamoxifen-induced iDKO and iCKO lungs. Principal component analysis separated iDKO from iCKO samples (**Figure 3A**), indicating broad genotype-dependent remodeling of the PH lung transcriptome. A global differentially expressed gene heatmap confirmed clear genotype-dependent clustering (**Figure 3B**). Volcano plot analysis showed that many of the most significantly reduced transcripts in iDKO lungs encoded cell-cycle and mitotic regulators, including Ccnb2, Cenpf, Ccna2, Cdk1, Esco2, and Top2a (**Figure 3C**). Hallmark GSEA confirmed strong negative enrichment of E2F targets, G2M checkpoint, and epithelial-mesenchymal transition programs in iDKO relative to iCKO lungs (**Figure 3D-F**). KEGG analysis of genes downregulated in iDKO identified cell cycle as the top enriched pathway, together with related mitotic programs and ECM-receptor interaction (**Figure 3H**). Heatmap shows that the core genes for E2F/G2M cell-cycle genes (Mki67, Top2a, Cdk1, Ccna2, Ccnb1, Ccnb2, Plk1, Cenpf, Birc5, Mcm2, Mcm6, and Pcna) and EMT/ECM-remodeling genes (Fn1, Col1a1, Col1a2, Col3a1, and Tnc) (**Figure 3I**). Together, these whole-lung data show that genetic E2F1 loss suppresses a proliferative, E2F/G2M-dominant,and matrix-remodeling component transcriptional program in *Egln1*-driven PH lungs.

**Figure 3.**
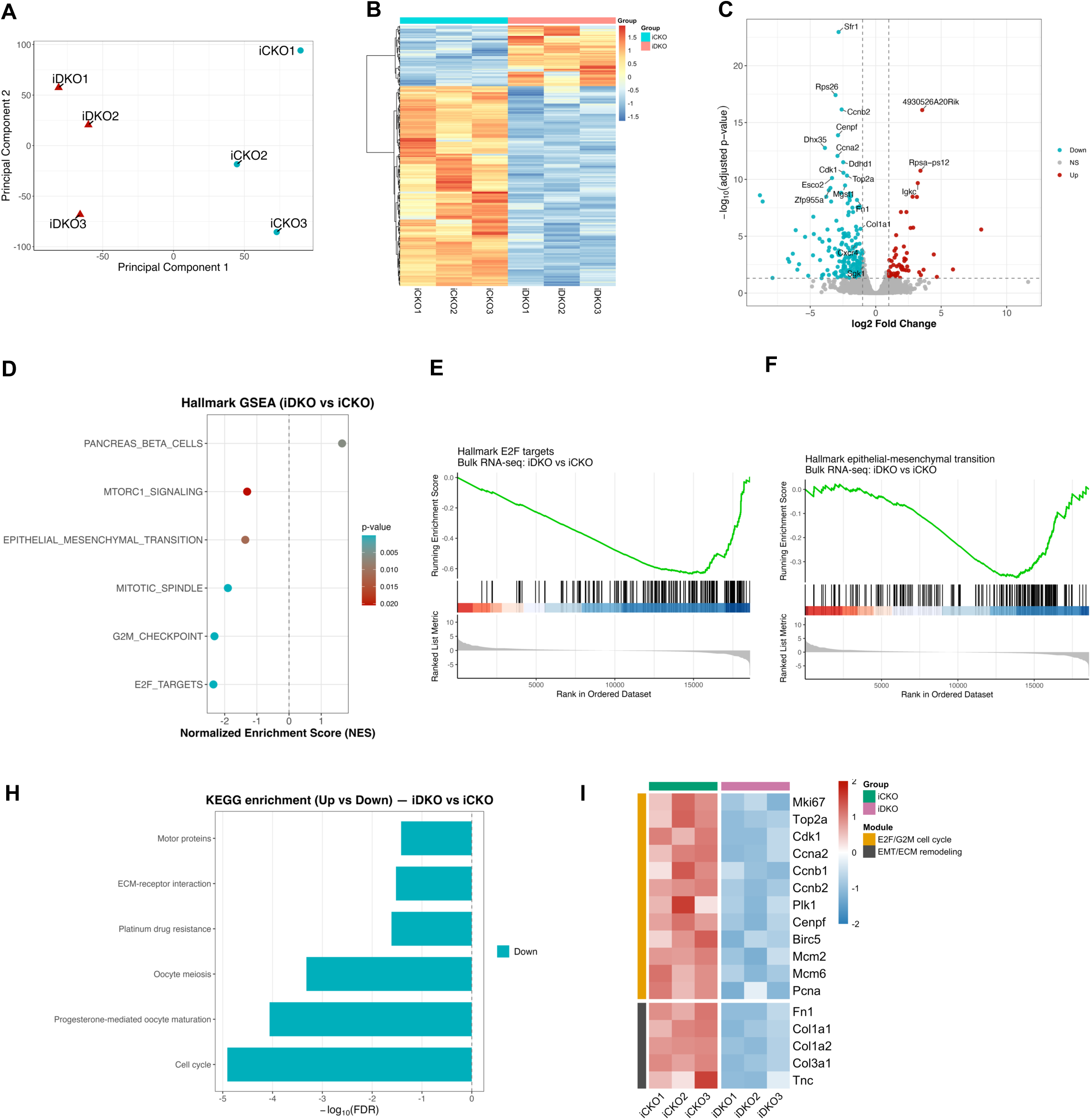
Whole-lung bulk RNA-seq shows that E2f1 deletion suppresses cell-cycle, mitotic, and EMT/ECM-remodeling programs in *Egln1*-driven PH lungs. (A) Principal component analysis (PCA) of whole-lung bulk RNA-seq from iCKO and iDKO mice (n = 4/group). Individual samples are labeled. (B) Heatmap of global differentially expressed genes in iDKO vs iCKO lungs. Rows represent genes and columns represent individual samples; values are shown as scaled expression. (C) Volcano plot of differential gene expression in iDKO vs iCKO lungs. Colored dots indicate significantly upregulated or downregulated genes, and gray dots indicate non-significant genes. Labeled downregulated genes include cell-cycle and mitotic regulators such as Ccnb2, Cenpf, Ccna2, Cdk1, Esco2, and Top2a. Differential expression was analyzed with DESeq2 and Benjamini-Hochberg FDR correction. (D-F) Hallmark GSEA of iDKO vs iCKO whole-lung bulk RNA-seq. (D) Dot plot of selected Hallmark pathways. Negative NES values indicate pathways suppressed in iDKO relative to iCKO. (E and F) Running enrichment plots for Hallmark E2F targets and epithelial-mesenchymal transition, respectively. (H) KEGG pathway enrichment analysis of genes downregulated in iDKO vs iCKO lungs, highlighting cell-cycle and mitotic programs. (I) Focused heatmap of predefined E2F/G2M cell-cycle and EMT/ECM-remodeling genes significantly reduced in iDKO lungs (adjusted P < 0.05 and log2FC < 0). Rows represent genes and columns represent individual samples; row annotations denote retained modules.

### Single-cell RNA-seq shows E2F1 loss attenuates pathological arterial reprogramming in *Egln1*-driven PH

To define cellular and endothelial programs associated with the hemodynamic rescue in iDKO mice, we performed whole-lung scRNA-seq from iCKO and iDKO lungs. After integration and annotation, 16,978 cells were retained, including 9,364 iCKO and 7,614 iDKO cells (**Supplemental Figure 2A**). Final annotation resolved 26 Seurat clusters into 31 cell populations encompassing epithelial, endothelial, mesenchymal, and immune lineages. (**Figure 4A and Supplemental Figure 2B**). Cell identities were validated using a 55-gene marker panel derived from the LungMAP CellCards framework^28^ (**Supplemental Figure 2C**). Cell composition and cell-type-resolved differential-expression analyses showed genotype-dependent remodeling of lung cell states, including changes in inflammatory, epithelial, mesenchymal, and endothelial populations (**Figure 4B-C and Supplemental Figure 2D**). These data indicate that E2F1 loss is associated with broad remodeling of the PH lung cell atlas, with a prominent effect on the endothelial compartment.

**Figure 4.**
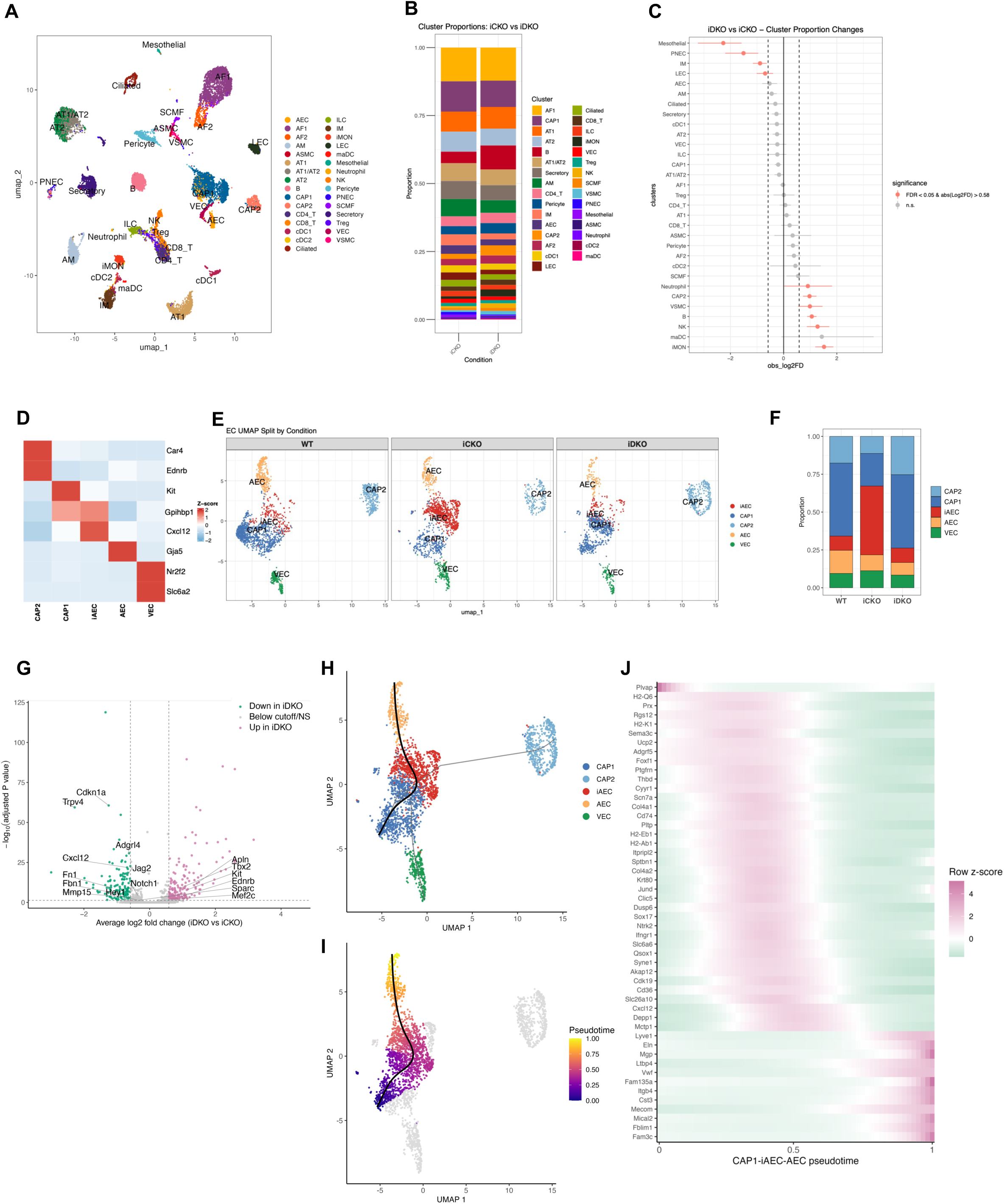
Single-cell RNA-seq demonstrates E2F1 loss attenuates pathological gCap (CAP1) to arterialization. (A) UMAP visualization of integrated whole-lung scRNA-seq from iCKO and iDKO mice. The final dataset contained 16,978 cells, including 9,364 iCKO and 7,614 iDKO cells. Major lung cell types are annotated, including immune, epithelial, mesenchymal, and endothelial compartments. (B) Stacked bar plot showing the proportion of major lung cell types in iCKO and iDKO lungs. (C) Differential cell proportion analysis using scProportionTest permutation testing. (D) EC subtype marker or annotation summary used to define CAP1, CAP2, iAEC, AEC, and VEC endothelial states. (E) Split UMAP visualizations showing the shift of gCap (CAP1) endothelial cells toward an arterial fate in iCKO lungs and its rescue in iDKO lungs. (F) Proportion of computationally subsetted endothelial subtypes across WT, iCKO, and iDKO lungs. (G) Volcano plot of all-EC differential expression in iDKO vs iCKO ECs. Differential expression was analyzed with Seurat FindMarkers using the Wilcoxon rank-sum test; significant genes were defined by adjusted P < 0.05 and |average log2FC| > log2(1.5) unless otherwise indicated. (H) Slingshot trajectory projected onto the endothelial UMAP. CAP1 was used as the root state, and CAP1-rooted branches included CAP1-iAEC-AEC, CAP1-iAEC-CAP2, and CAP1-VEC trajectories. (I) Pseudotime projection for the CAP1-iAEC-AEC trajectory. Color indicates scaled pseudotime from early to late positions along the trajectory. (J) tradeSeq heatmap of dynamic genes along CAP1-iAEC-AEC pseudotime. Genes shown were significant by associationTest after fitGAM modeling (Benjamini-Hochberg FDR < 0.05).

Because the genetic lesion and rescue phenotype were endothelial-centered, we next computationally subsetted ECs from the whole-lung scRNA-seq dataset. EC analysis resolved five endothelial states: AEC, CAP1, CAP2, VEC, and intermediate AEC (iAEC) (**Supplemental Figure 3A**). Compared with WT and iCKO controls, iDKO lungs showed reduced iAEC accumulation and a more normalized CAP1/CAP2 distribution, indicating that E2F1 loss blunts the maladaptive endothelial-state transition that emerges in *Egln1*-driven PH (**Figure 4D-F**). These subtype assignments were used for trajectory and all-EC differential-expression analyses.

All-EC differential expression visualized by volcano plot further defined the E2F1-dependent endothelial program (**Figure 4G and Supplemental Figure 3B**). Using adjusted P < 0.05 and |average log2FC| > 0.585, 124 genes were reduced and 166 genes were increased in iDKO ECs relative to iCKO ECs. Genes reduced in iDKO included Cdkn1a, Trpv4, Cxcl12, Jag2, Adgrl4, and EMT/ECM-remodeling genes Fn1, Fbn1, and Mmp15. Notch1, Hey1, and Sparc were also significantly reduced, although their fold changes were below the volcano cutoff. In contrast, CAP2 markers Apln, Ednrb, Tbx2 and CAP1 marker Kit and Mef2c were increased in iDKO ECs. Together, these single-cell data show that E2F1 loss shifts ECs away from a arterial remodeling state while attenuating ECM-associated endothelial activation.

Trajectory analysis further supported this shift away from pathological endothelial remodeling. Slingshot pseudotime analysis of computationally subsetted ECs identified three CAP1-rooted branches, including CAP1-to-iAEC-to-CAP2, CAP1-to-iAEC-to-AEC, and CAP1-to-VEC trajectories. We focused on the CAP1-to-iAEC-to-AEC branch as the disease-relevant arterializing trajectory (**Figure 4H**). Compared with iCKO ECs, iDKO ECs were shifted toward earlier positions along this branch, with lower median scaled pseudotime and a smaller fraction of late-pseudotime cells (**Figure 4I**). To identify genes dynamically regulated along this trajectory, we applied tradeSeq using fitGAM followed by associationTest. Among 5,998 expressed genes modeled, 1,545 genes were significantly associated with CAP1-to-iAEC-to-AEC pseudotime (FDR < 0.05), including extracellular matrix and vascular-remodeling genes Col4a1, Col4a2, Eln, Mgp, Thbd, Ltbp4, Vwf, and Cxcl12, together with Sox17 and other endothelial-state regulators (**Figure 4J**). Several volcano-highlighted genes were also dynamic along AEC pseudotime, including Cxcl12, Adgrl4, Cdkn1a, Gja5, Jag2, Kit, Mmp15, Fbn1, Trpv4, Fn1, Ednrb, Hey1, Mef2c, and Tbx2. Finally, tradeSeq diffEndTest identified 2,070 genes with significant endpoint differences among the terminal CAP2, AEC, and VEC branches (global FDR < 0.05), supporting branch-specific EC terminal programs. Together, these trajectory-based analyses indicate that E2F1 loss reduces progression of ECs into a remodeled AEC state and attenuates matrix-remodeling and arterializing gene programs coupled to this transition.

### E2F1 loss disrupts endothelial cell communication hub

To define how E2F1 loss reshapes intercellular signaling within the PH lung, we inferred cell–cell communication across all 31 annotated cell populations in the iCKO and iDKO scRNA-seq datasets using CellChat with the curated mouse ligand–receptor database. E2F1 loss produced a global reduction of intercellular signaling: the number of inferred interactions decreased modestly, total interaction strength fell by ∼25% (**Figure 5A**). Differential network analysis localized this loss to the endothelial compartments (**Supplemental Figure 4A**). Ranking pathways by information flow identified a coherent set of programs significantly reduced in iDKO, led by extracellular-matrix and basement-membrane signaling (COLLAGEN, LAMININ, FN1), angiogenic and arterial programs (VEGF, ANGPTL, NOTCH), and additional remodeling-associated pathways (THBS, SEMA3, SEMA4, JAM) (**Figure 5B and Supplemental Figure 4B**). Several endothelial pathways were extinguished entirely in iDKO, including VWF, VTN, SPP1, VCAM, and PERIOSTIN. This coordinated reduction of matrix, angiogenic/arterial, and endothelial-activation signaling parallels the bulk and single-cell transcriptional signatures and indicates that E2F1 loss protects the lung from a matrix-remodeling, arterializing communication state.

**Figure 5.**
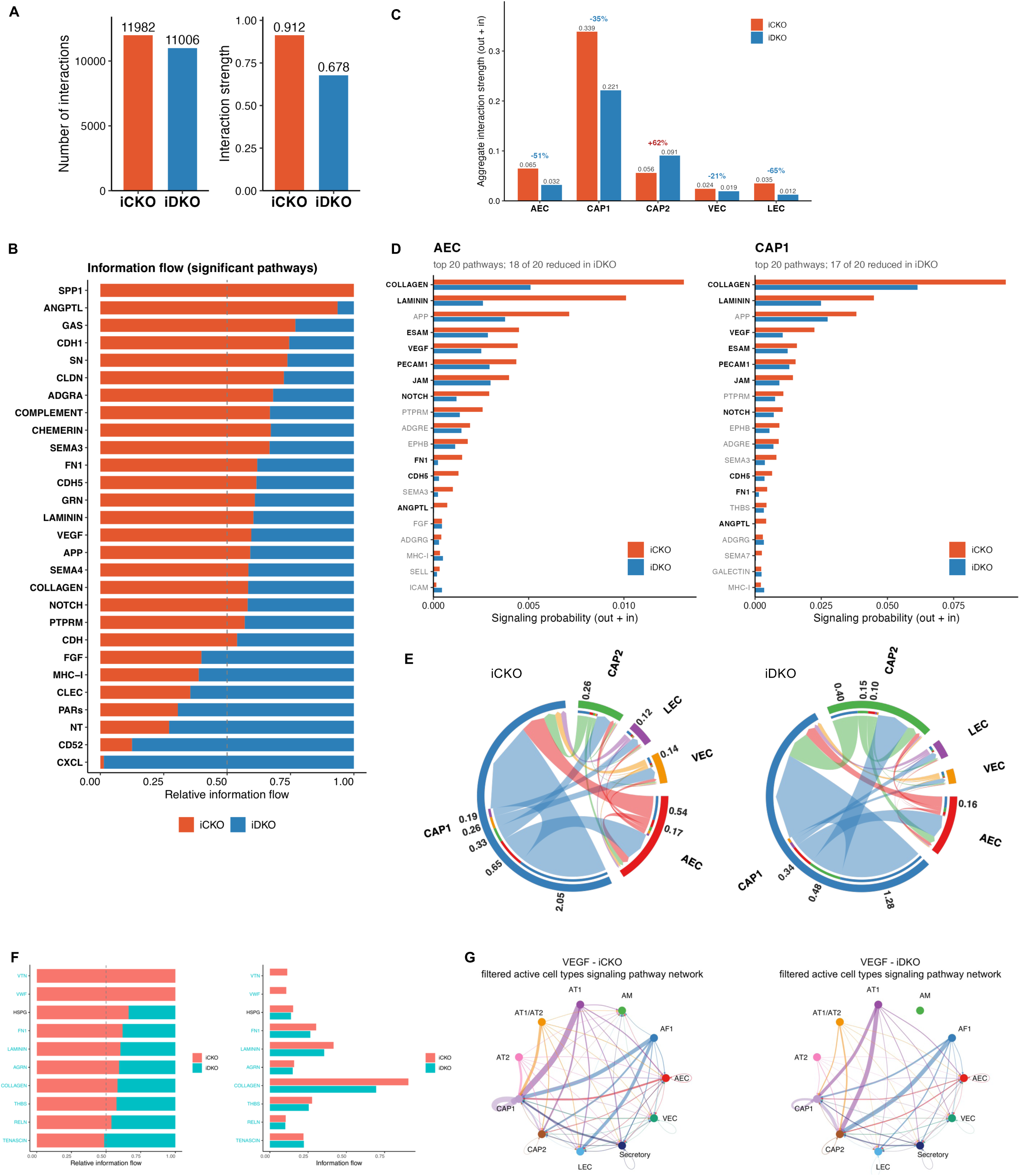
E2F1 loss disrupts an arterial endothelial cell–cell communication hub. Cell-cell communication was inferred from iCKO and iDKO whole-lung scRNA-seq datasets using CellChat v2.1.2 with the mouse ligand-receptor database. Analyses used log-normalized RNA assay values, triMean communication probabilities, population-size correction, and min.cells = 10; n = 9,364 iCKO cells and 7,614 iDKO cells across 31 annotated cell populations. (A) Total inferred interaction number and aggregate interaction strength per condition. E2F1 loss modestly reduced interaction number and produced a larger reduction in total interaction strength in iDKO lungs. (B) Relative information flow of significantly altered signaling pathways between iCKO and iDKO. Pathways preferentially enriched in iCKO included extracellular-matrix and basement-membrane programs that were reduced in iDKO. (C) Aggregate endothelial signaling strength, calculated as summed outgoing plus incoming interaction strength, across endothelial subtypes. AEC and CAP1 signaling were reduced in iDKO, whereas CAP2 showed increased signaling strength. (D) Top pathway-level signaling contributions for AEC and CAP1, showing coordinated reduction of matrix, arterial/angiogenic, and endothelial-junction signaling modules in iDKO. (E) NOTCH signaling chord diagrams for iCKO and iDKO, showing reduced endothelial NOTCH communication after E2F1 loss. (F) ECM-Receptor information flow comparing iCKO and iDKO. COLLAGEN remained the dominant ECM pathway but was reduced in iDKO, together with decreases in LAMININ, FN1, THBS, HSPG, and AGRN; VTN and VWF signaling were lost. (G) VEGF signaling networks in iCKO and iDKO after filtering to active interacting cell types. VEGF signaling was reduced and redistributed in iDKO, with diminished CAP1-centered communication and relative preservation or rerouting toward CAP2.

Because the genetic lesion, the rescue phenotype, and the transcriptional program were all endothelial-centered, we quantified the aggregate signaling activity of each EC subpopulations. The arterial endothelial cell (AEC) and CAP1 were among the two most strongly affected populations (**Figure 5C**). Notably, CAP2 capillaries instead increased their aggregate signaling (**Figure 5C**). Decomposing AEC and CAP1 signaling by pathway showed that every AEC-associated pathway decreased in iDKO, spanning three coherent modules: basement-membrane matrix (COLLAGEN, LAMININ, FN1), arterial/angiogenic signaling (VEGF, NOTCH, ANGPTL), and endothelial cell–junction and adhesion programs, including VE-cadherin, PECAM1, ESAM, and JAM (**Figure 5D**). At the communication level, therefore, E2F1 loss reduced arterial-endothelial identity, including NOTCH signaling, a canonical arterial-specification program17. Total NOTCH signaling fell ∼28% in iDKO and was overwhelmingly endothelial-driven, with all Notch ligands (*Jag2*, *Dll4*, *Jag1*, *Dll1*) originating from endothelial cells. The dominant ligand–receptor pair, *Jag2*–*Notch1*, was the most reduced, consistent with the reduced *Jag2* expression in iDKO ECs, and the AEC contribution as a NOTCH source dropped ∼67% — the largest proportional loss among senders (**Figure 5E** and **Supplemental Figure 4C**). These data place E2F1 upstream of arterial-specification Notch signaling in the pulmonary endothelium.

Extracellular-matrix signaling was the single largest communication loss in iDKO (**Figure 5F**). COLLAGEN, LAMININ, and FN1 accounted for essentially the entire effect, with vitronectin and von Willebrand factor signaling abolished. The VEGF angiogenic axis behaved similarly: *Vegfa*–*Vegfr2*, *Vegfa*–*Vegfr1*, and *Vegfa*–*Vegfr1r2* signaling all decreased^36^, the dominant capillary receiver (CAP1) collapsed by 61%, and the residual signal re-routed toward CAP2 (**Figure 5G** and **Supplemental 3D**). Together, these analyses show that E2F1 loss selectively disrupts an arterial/capillary endothelial communication hub, inhibiting Notch arterial-specification, VEGF/ANGPTL angiogenic, and basement-membrane matrix signaling. This intercellular signature provides an independent, network-level correlate of the E2F1-dependent endothelial arterial-remodeling program.

### E2F1 overexpression in HLMVECs drives cell-cycle and Notch/arterial programs

To dissect cell-autonomous consequences of E2F1 gain-of-function, we transduced HLMVECs with AdvE2F1 or control adenovirus. qPCR and immunoblot analyses confirmed robust E2F1 overexpression (**Figure 6A and 6B**), and BrdU/DAPI imaging showed increased BrdU incorporation, indicative of increased proliferation in AdvE2F1-transduced ECs (**Figure 6C**). Bulk RNA-seq of AdvE2F1 vs AdvCTL HLMVECs revealed clear separation between groups by PCA (**Figure 6D**). Volcano plot analysis highlighted significant induction of E2F/cell-cycle genes and Notch/arterial-associated genes, including E2F1, E2F2, PCNA, MCM3, MCM4, MCM7, CCNE1, CCNE2, CDC25A, CDKN1A, HEY1, HEY2, HES6, and NOTCH3 (**Figure 6E**). A global DEG heatmap confirmed broad transcriptional remodeling (**Figure 6F**). Hallmark and KEGG pathway analyses further showed enrichment of proliferative and signaling programs, with KEGG upregulated pathways including DNA replication, cell cycle, p53 signaling, and Hippo signaling, and downregulated pathways including the TCA cycle and carbon metabolism (**Figure 6G, 6H, and 6I**). To focus the main mechanistic signature, we generated a curated heatmap restricted to statistically significant upregulated genes (adjusted P < 0.05), which contained 28 E2F/cell-cycle genes and 11 Notch/arterial genes (**Figure 6J**). At the protein level, AdvE2F1 increased arterial EC marker SOX17 abundance (**Figure 6K**) and reduced BMPR2 under basal and BMP9-stimulated conditions (**Supplemental Figure 4**). Together, these data indicate that E2F1 is sufficient to activate proliferative and Notch/arterial-associated transcriptional programs while impairing BMPR2 protein responses in pulmonary ECs.

**Figure 6.**
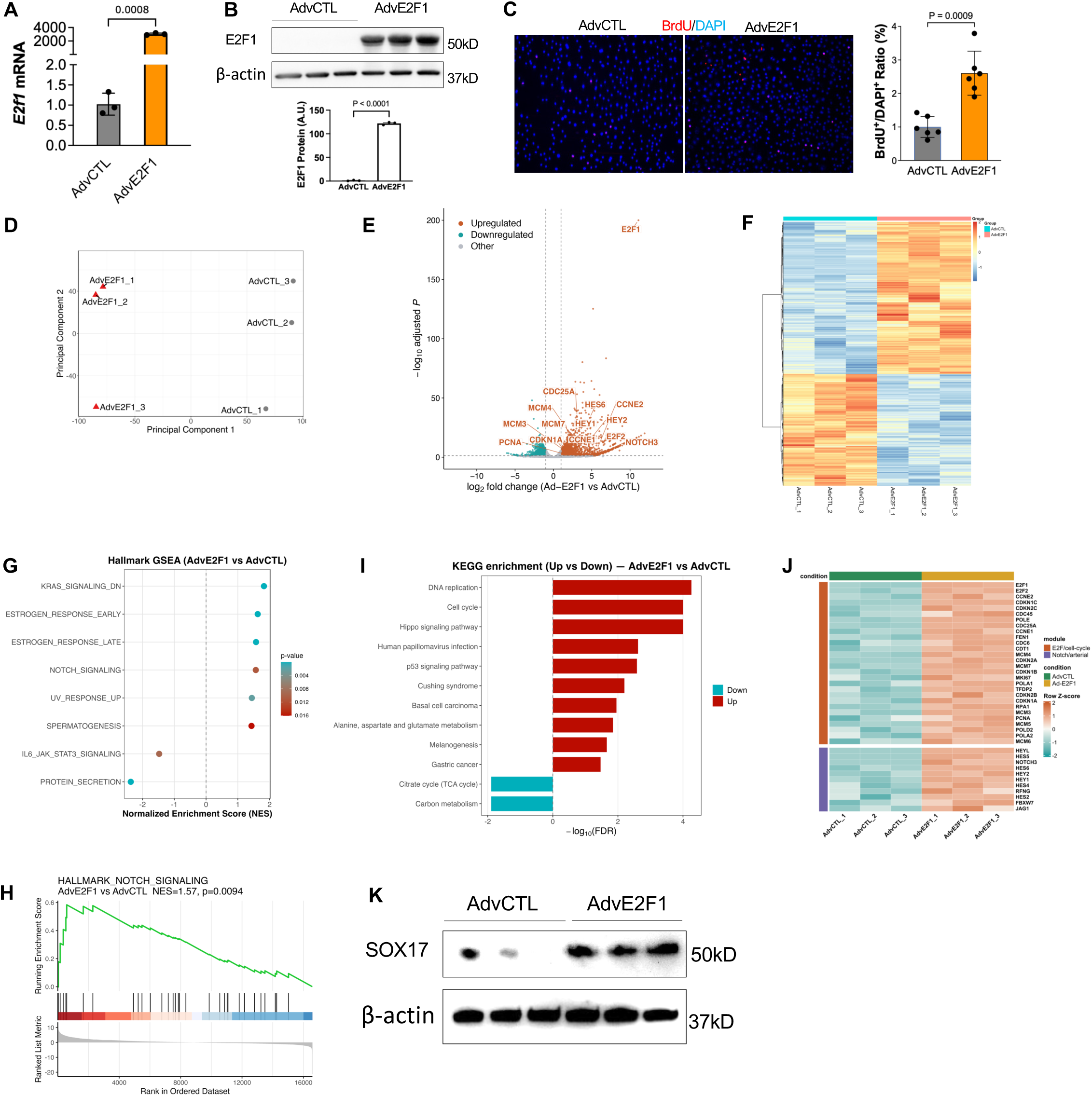
E2F1 overexpression in HLMVECs drives proliferative and arterial programs. (A) qPCR analysis of E2F1 mRNA expression in AdvCTL- and AdvE2F1-transduced HLMVECs. Data are normalized to AdvCTL controls. (B) Representative immunoblot of E2F1 protein in HLMVECs transduced with AdvCTL or AdvE2F1; β-actin served as loading control. (C) Representative BrdU/DAPI immunofluorescence images and quantification of BrdU-positive nuclei in AdvCTL- and AdvE2F1-transduced HLMVECs. Scale bar = 50 µm. (D) PCA of bulk RNA-seq data from AdvCTL- and AdvE2F1-transduced HLMVECs (n = 3/group). (E) Focused volcano plot of bulk RNA-seq differential expression in AdvE2F1 vs AdvCTL HLMVECs. Labeled significant genes include E2F and Notch targets. Differential expression was analyzed with DESeq2 and Benjamini-Hochberg FDR correction. (F) Heatmap of global differentially expressed genes in AdvE2F1 vs AdvCTL HLMVECs. Rows represent genes and columns represent samples; color indicates scaled normalized expression. (G) Hallmark GSEA dot plot for AdvE2F1 vs AdvCTL HLMVECs. Positive NES values indicate enrichment in AdvE2F1 cells. (H) Running enrichment plot for Hallmark NOTCH signaling upregulated in AdvE2F1 HLMVECs. (I) KEGG enrichment analysis of upregulated and downregulated genes. (J) Focused heatmap of statistically significant upregulated genes (adjusted P < 0.05) in E2F/cell-cycle and Notch/arterial modules. (K) Representative immunoblot of SOX17 protein in AdvCTL- and AdvE2F1-transduced HLMVECs. For qPCR, BrdU quantification, and immunoblot densitometry, individual points indicate independent biological replicates and two-group comparisons used unpaired two-tailed t tests unless otherwise indicated.

### Pharmacological E2F inhibition attenuates *Egln1*-driven PH

We next tested whether pharmacological E2F inhibition could suppress endothelial proliferation and mitigate Egln1-driven PH. HLM006474 was administered to tamoxifen-induced *Egln1^Cdh5CreERT^*^2^ mice during the established PH window (**Figure 7A**). HLM006474 reduced RVSP and pulmonary vascular remodeling compared with vehicle-treated controls, with a more modest effect on RV hypertrophy (**Figure 7B-C**). Vascular remodeling analysis showed reduced distal pulmonary arterial wall thickening and luminal narrowing after HLM006474 treatment (**Figure 7D-E**), and α-SMA staining showed reduced α-SMA^+^ distal pulmonary arterial muscularization (**Figure 7F-G**). In parallel in vitro assays, HLM006474 suppressed VEGF-A- and hypoxia-induced BrdU incorporation in HLMVECs (**Figure 7H-I**). These data indicate that pharmacological E2F inhibition attenuates Egln1-driven PH in mice.

**Figure 7.**
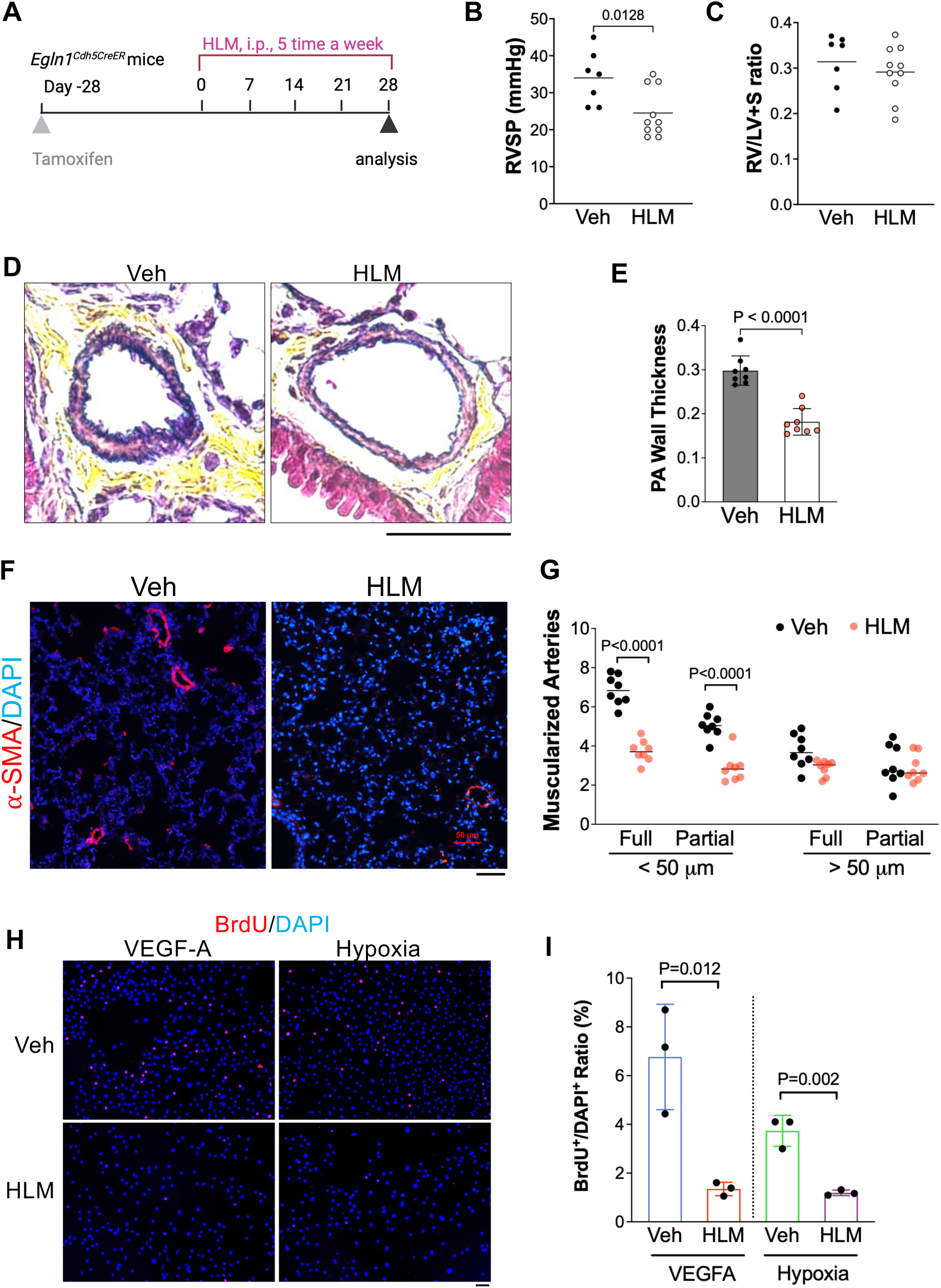
Pharmacological inhibition of E2F suppresses endothelial proliferation and attenuates *Egln1*-driven PH. **(A)** Experimental timeline for HLM006474 treatment in tamoxifen-induced *Egln1^Cdh5CreERT^*^2^ mice. Tamoxifen was administered before HLM006474 treatment, and analysis was performed after the treatment period. (B) HLM006474 treatment reduced RVSP compared with vehicle-treated controls. (C) Right ventricular hypertrophy was quantified by Fulton index, RV/(LV+S). (D) Representative Russell-Movat pentachrome-stained distal pulmonary arteries showing vascular wall thickening in vehicle-treated Egln1Cdh5CreERT2 mice, with reduced remodeling after HLM006474 treatment. Scale bar = 50 µm. (E) Quantification of pulmonary vascular wall thickening. (F) Representative α-SMA immunofluorescence images of distal pulmonary arteries showing reduced smooth muscle coverage after HLM006474 treatment. Scale bar = 50 µm. (G) Quantification of α-SMA-positive distal pulmonary arterial muscularization. (H and I) Representative BrdU/DAPI immunofluorescence images and quantification from HLMVECs treated with vehicle or HLM006474 under VEGF-A stimulation or hypoxia. Scale bar = 50 µm. HLM006474 suppressed VEGF-A- and hypoxia-induced BrdU incorporation. Individual points indicate biological replicates; n values are shown in each graph. Two-group comparisons used unpaired two-tailed t tests, and multi-condition BrdU analyses used one-way ANOVA with post hoc correction as appropriate.

### HLM006474 attenuates monocrotaline-induced PH in rats

We next validated the therapeutic efficacy of HLM006474 in the severe PH, MCT rat model. In the prevention protocol, HLM006474 was administered after MCT exposure and significantly reduced RVSP and RV/LV+S ratio, improved survival, and attenuated vascular remodeling by Russell-Movat pentachrome (**Figure 8A-F**). In a reversal protocol in which HLM006474 was started after PH was established, HLM006474 also reduced RVSP and pulmonary vascular remodeling, although RV/LV+S ratio was not significantly changed (**Figure 8G-M**). These pharmacological data support a model in which E2F1 drives PH progression through coordinated effects on cell-cycle activation, Notch-associated arterial programming, and pulmonary vascular remodeling (**Figure 8N**).

**Figure 8.**
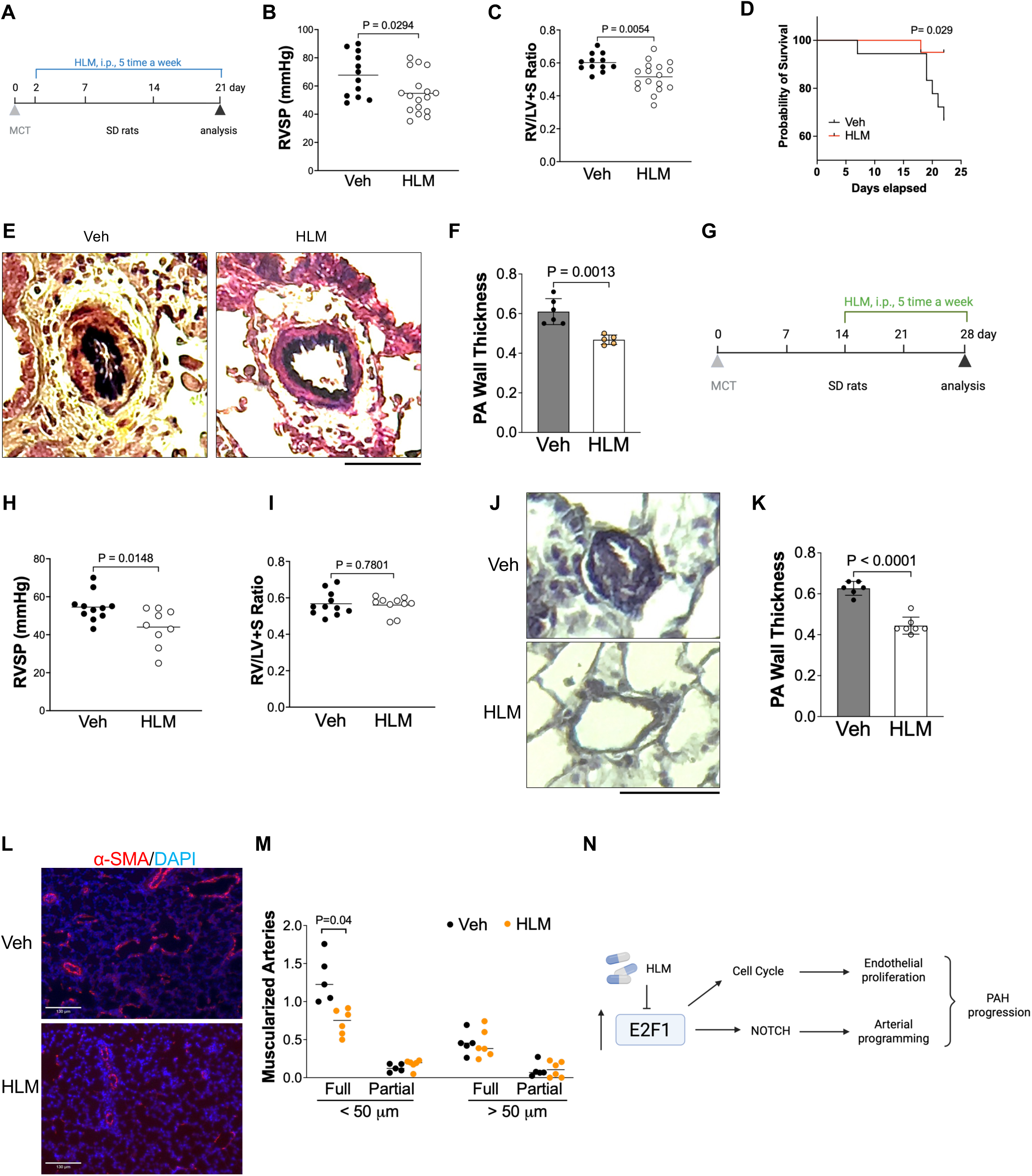
Pharmacological inhibition of E2F1 attenuates and reverses MCT-induced PH in rats. (A) Prevention-protocol timeline for HLM006474 treatment in monocrotaline (MCT)-induced PH rats. HLM006474 treatment was initiated after MCT exposure and continued until terminal analysis. (B) Prevention-protocol RVSP measurements showing reduced PH severity with HLM006474. (C) Right ventricular hypertrophy quantified by RV/(LV+S). (D) Survival analysis of vehicle- and HLM006474-treated MCT rats. (E and F) Representative Russell-Movat pentachrome staining and quantification of pulmonary vascular remodeling in vehicle- and HLM006474-treated MCT rats. Scale bar = 50 µm. (G) Reversal-protocol timeline in MCT rats, in which HLM006474 treatment was initiated after PH was established. (H and I) Reversal-protocol RVSP and RV/(LV+S) measurements; HLM006474 reduced RVSP, whereas RV hypertrophy was not significantly changed. (J and K) Representative Russell-Movat pentachrome staining and quantification of pulmonary vascular remodeling in the reversal protocol. Scale bar = 50 µm. (L and M) Representative α-SMA/DAPI immunofluorescence images and quantification of distal pulmonary arterial muscularization in the reversal protocol. Scale bar = 50 µm. (N) Working model. E2F1 promotes endothelial proliferation and arterial programming to drive PAH progression; HLM006474 inhibits E2F activity and attenuates these remodeling programs. Individual points indicate animals, and n per group is shown in each graph. Two-group comparisons used unpaired two-tailed t tests, multi-group comparisons used ANOVA with post hoc correction where applicable, and survival was analyzed by log-rank test.

## Discussion

The principal novelty of this study is two-fold. First, we identify E2F1 as a previously underappreciated driver of pulmonary vascular remodeling and PAH, rather than simply a downstream marker of cell-cycle entry. Second, we define a new endothelial mechanism in which E2F1 promotes Notch-associated arterial programming and AEC trajectory progression, linking proliferative pressure to pathological endothelial-state remodeling. Five lines of evidence support this conclusion. First, E2F1 expression and nuclear protein are elevated in human PAH samples and in multiple preclinical PH models. Second, loss of E2F1 in the *Egln1*-driven model rescues key hemodynamic and structural features of PH. Third, whole-lung bulk RNA-seq and single-cell profiling show that E2F1 loss suppresses E2F/G2M transcriptional programs and attenuates maladaptive endothelial reprogramming, including induced arterial endothelial cell states and matrix-remodeling programs. Fourth, gain-of-function studies in HLMVECs show that E2F1 is sufficient to promote endothelial proliferation, activate E2F/cell-cycle and Notch/arterial-associated transcriptional programs, increase SOX17 abundance, and suppress BMPR2 protein responses. Fifth, pharmacological pan-E2F inhibition with HLM006474 attenuates PH in both *Egln1*-driven and MCT models. Together, these data position E2F1 as a tractable transcriptional hub in PAH pathogenesis.

Our findings reposition E2F1 from a generic cell-cycle effector to a disease-specific endothelial remodeling factor in PAH. Prior work linked endothelial SOX17, a transcription factor encoded by a PAH risk locus,^7^ to E2F1 induction and SOX17-deficiency-induced PH.^18^ Earlier work in PASMCs identified an NHE1-p27-E2F1 axis sufficient to drive PASMC proliferation,^17^ and a separate body of work demonstrated that decreased Notch2 expression in PAECs enhances E2F1 binding to the NOTCH1 promoter and supports a proliferative, apoptosis-resistant phenotype.^37^ Our data extend these observations by showing that E2F1 is required in vivo for severe pulmonary vascular remodeling and is sufficient in pulmonary ECs to activate an arterializing, Notch-associated transcriptional state. Thus, persistent endothelial E2F1 activation integrates cell-cycle entry, BMPR2 suppression, and ARF-p53-p21 surveillance with a maladaptive arterial-programming response that characterizes PAH endothelium.^38,39^

A complementary mechanistic finding is the suppression of BMPR2 by E2F1 in HLMVECs. Heterozygous BMPR2 loss-of-function is the most penetrant genetic cause of heritable PAH, and acquired BMPR2 deficiency is a near-universal feature of IPAH endothelium.^5,6^ Selective reactivation of endothelial BMPR2 reverses experimental PAH,^40^ defining BMPR2 restoration as an attractive therapeutic node. Our data suggest that pharmacological E2F1 inhibition may indirectly restore BMPR2 signaling and thereby converge with BMPR2-based strategies on a shared endothelial homeostatic program.

The second major mechanistic advance is the identification of E2F1 as a regulator of endothelial arterial programming. Recent lineage-tracing and single-cell transcriptomics demonstrate that general capillary ECs transit toward an arterial identity in PH lungs through a HIF-2α/Notch4 axis,^19^ producing an induced arterial EC population. Our scRNA-seq data place E2F1 within this circuit: loss of E2F1 in iDKO mice contracts the AEC fraction, shifts ECs toward earlier CAP1-iAEC-AEC pseudotime positions, and reduces expression of PH-remodeling genes including Cdkn1a, Trpv4, Cxcl12, Jag2, Adgrl4, Fn1, Fbn1, and Mmp15. Conversely, gain of E2F1 in HLMVECs induces significant E2F/cell-cycle and Notch/arterial-associated genes, including HEY1, HEY2, HES6, and NOTCH3. These reciprocal loss- and gain-of-function data suggest that E2F1 is not merely permissive for proliferation but actively helps specify the maladaptive arterial/iAEC program that accompanies PH vascular remodeling.

The induction of an ARF-p53-p21 senescence signature in E2F1-overexpressing HLMVECs is, on its face, counterintuitive: E2F1 is canonically pro-proliferative. However, in the setting of sustained, supraphysiological E2F1 activity, the ARF-p53-p21 arm is engaged as a tumor-suppressive surveillance program,^14,15^ producing a “proliferation-poised but senescence-permissive” state. This state appears to be precisely what is observed in PAH endothelium: pulmonary ECs in PAH lungs are simultaneously hyperproliferative and enriched for senescence markers including p21 and p16, and senescent ECs have been shown to drive PASMC proliferation through Notch signaling.^38,39,41^ Our iDKO scRNA-seq data—specifically the normalization of *Cdkn1a* (p21) in iAEC populations—provide an *in vivo* genetic test of this model: endothelial E2F1 is required for the accumulation of p21-positive senescence-marked ECs in PH lungs.

From a translational standpoint, our results support E2F inhibition as a candidate disease-modifying strategy in PAH. HLM006474 attenuated PH in both an *Egln1*-driven endothelial model and the MCT rat model, including a reversal protocol in rats (**Figure 8**). In vitro, HLM006474 directly suppressed VEGF-A- or hypoxia-induced EC proliferation. Because HLM006474 is a pan-E2F inhibitor, future work will need to determine whether selective E2F1 inhibition is sufficient and whether combination strategies with sotatercept, HIF-2α antagonism, BMPR2 restoration, or Notch-pathway modulation provide additive benefit.

This study has several limitations. First, iDKO mice carry global (whole-body) E2f1 deletion on an endothelial *Egln1*-knockout background, rather than endothelial-specific E2f1 deletion. Although our experiments demonstrate that endothelial E2F1 is required for PH development, the constitutive loss of E2f1 in all cell types leaves open the possibility that E2F1 in smooth muscle cells, hematopoietic cells may also contribute to the phenotype. Future studies with conditional E2f1 flox alleles crossed with endothelial-specific Cre drivers (e.g., Cdh5-CreERT2) will be needed to formally establish endothelial cell-autonomy. Second, HLM006474 is a pan-E2F inhibitor and its effects on E2F members cannot be excluded; isoform-selective compounds and structural data on E2F1-DNA-binding inhibition will be required to refine drug development. Third, while our reversal protocol in MCT rats demonstrates therapeutic potential, further reversal studies in Sugen/hypoxia rats are a logical next step. Finally, although our human data show E2F1 elevation in IPAH ECs, biomarker studies linking pulmonary or circulating E2F1 activity readouts to PAH severity, progression, or therapeutic response remain to be performed.

In summary, E2F1 is a novel transcriptional driver of pulmonary vascular remodeling in PAH. Beyond its canonical role in cell-cycle control, E2F1 regulates an endothelial arterial-programming mechanism marked by Notch-associated signaling, AEC trajectory progression. Genetic loss or pharmacological inhibition of E2F activity mitigates experimental PH across complementary models, supporting E2F1 as a candidate therapeutic node for PAH and positioning E2F1-dependent arterial programming as a mechanistic signature of disease-modifying response.

## Sources of Funding

This work was supported in part by National Institutes of Health grants R01HL158596, R01HL162794, R01HL169509, R01HL170096, American Heart Association Career Development Award 20CDA35310084, and institutional support to Z.D.

## Disclosures

The authors declare no competing financial interests.

## Acknowledgments

None

## Supplemental Figure legends

**Supplemental Figure 1. Egln1Tie2Cre lungs show induction of an arterial/endothelial remodeling gene program.** Heatmap showing row-scaled expression of selected arterial, endothelial activation, and extracellular-matrix remodeling genes in WT and Egln1Tie2Cre (CKO) mouse lungs (n = 3/group). Each column represents an individual mouse sample. Color indicates row z score, with blue indicating lower expression and red indicating higher expression.

**Supplemental Figure 2. Whole-lung scRNA-seq annotation and cell-type-resolved differential expression in iCKO and iDKO lungs.** (A) UMAP visualization of integrated whole-lung scRNA-seq data showing initial clustering/automated annotation structure and final manually curated cell-type annotations. Final annotations resolved 16,978 cells into 31 epithelial, endothelial, mesenchymal, and immune populations. (B) Sankey plot mapping Seurat clusters to final cell-type annotations. Connections represent annotation contributions comprising >5% of a given cluster. (C) Dot plot showing expression of 55 LungMAP CellCards marker genes across final annotated cell populations, with iCKO and iDKO cells pooled. Dot size indicates the percentage of cells expressing each marker, and color indicates scaled average expression. (D) Jitter plot of cell-type-specific differential gene expression comparing iDKO vs iCKO using Seurat Wilcoxon testing. Each point represents a gene within a cell type, plotted by average log2 fold change. Gray points indicate non-significant genes; colored points indicate significant genes defined by FDR < 0.05 and |average log2FC| > log2(1.5).

**Supplemental Figure 3. Endothelial subclustering and arterial gene expression across WT, iCKO, and iDKO lungs**. (A) UMAP of computationally subsetted endothelial cells colored by Seurat cluster, showing endothelial substructure used to define AEC, CAP1, CAP2, VEC, and intermediate AEC (iAEC) states. (B) Endothelial UMAP colored by condition, showing the distribution of WT, iCKO, and iDKO ECs. (C) Heatmap of selected endothelial arterialization, capillary-state, Notch, matrix-remodeling, and injury-associated genes across WT, iCKO, and iDKO endothelial cells. Columns represent individual cells grouped by condition; rows represent selected genes. Color indicates scaled expression, with blue indicating lower expression and red indicating higher expression.

**Supplemental Figure 4. CellChat analysis of altered intercellular communication in iDKO vs iCKO lungs.** CellChat analysis used n = 9,364 iCKO and 7,614 iDKO cells, the mouse ligand-receptor database, triMean communication probabilities, population-size correction, and min.cells = 10. (A) Differential CellChat interaction heatmaps showing changes in inferred interaction number and interaction strength between annotated lung cell populations in iDKO relative to iCKO. Red indicates increased interactions in iDKO, whereas blue indicates decreased interactions. (B) RankNet information-flow analysis comparing signaling pathways between iCKO and iDKO lungs. Left, relative information flow; right, absolute information flow. E2F1 loss reduces multiple matrix, angiogenic, and endothelial-remodeling pathways, including COLLAGEN, LAMININ, FN1, VEGF, ANGPTL, and NOTCH. (C) NOTCH ligand-receptor pair contributions comparing iCKO and iDKO. Total NOTCH signaling is reduced in iDKO, with the dominant Jag2-Notch1 interaction decreased. (D) VEGF ligand-receptor pair contributions comparing iCKO and iDKO. Total VEGF signaling is reduced in iDKO. Bars indicate summed CellChat communication probability for each ligand-receptor pair.

**Supplemental Figure 5. E2F1 overexpression reduces BMPR2 signaling. HLMVECs** were transduced with control adenovirus (AdvCTL) or E2F1-expressing adenovirus (AdvE2F1) and analyzed under basal conditions or after BMP9 stimulation. Representative immunoblots show BMPR2 protein abundance in AdvCTL- and AdvE2F1-transduced cells. E2F1 overexpression reduced BMPR2 protein levels and attenuated BMP9-induced signaling compared with AdvCTL controls. β-actin or total protein served as loading control; where densitometry is shown, values were normalized to the loading control and then to the corresponding AdvCTL condition.

## Notes

### Competing Interest Statement

The authors have declared no competing interest.

